# Expression of the C-type lectin receptor CD205 on B cells mediates HIV-1 binding and *trans* infection of CD4^+^ T cells

**DOI:** 10.64898/2026.08.28.747757

**Authors:** Abigail D. Gerberick, Allison E. DePuyt, Peter E.J. Shoucair, Robbie B. Mailliard, Simon C. Watkins, Nicolas Sluis-Cremer, Charles R. Rinaldo

## Abstract

Antigen presenting cells (APC) can bind HIV-1 and subsequently *trans* infect CD4^+^ T cells. In comparison to direct (*cis*) infection of CD4^+^ T cells by free virus, APC-mediated HIV-1 *trans* infection is significantly more efficient and requires lower virus titers. As such, B cell-mediated HIV-1 *trans* infection of CD4^+^ T cells, particularly in secondary lymphoid organs (SLO) where B and CD4^+^ T cells interact frequently in and around B cell follicles, represents an efficient pathway for establishing and maintaining the latent HIV-1 reservoir. The molecular events involved in HIV-1 binding to B cells and transfer to CD4^+^ T cells are poorly understood. B cells are exposed to various activation signals in SLO including CD40 ligand (CD40L), interleukin-4 (IL-4), interferon-γ (IFN-γ), and B cell activating factor (BAFF). Here, we treated B cells with these different signals, or combination of signals, to identify those that facilitate HIV-1 binding to B cells and *trans*-infection of CD4^+^ T cells, and the mechanisms involved. We found that CD40L/IL-4 stimulated B cells are highly efficient mediators of HIV-1 *trans* infection of CD4^+^ T cells due to their enhanced capacity to bind HIV-1. Single cell RNA sequencing of differentially stimulated B cell populations revealed that CD40L/IL-4 stimulation significantly induced expression of the C-type lectin CD205. Confocal microscopy revealed that HIV-1 and CD205 co-localized on CD40L/IL-4 stimulated B cells, and antibody blocking of CD205 on these cells significantly reduced HIV-1 binding. Taken together, this study identifies CD205 as a critical receptor on B cells that facilitates HIV-1 binding and the transfer of virus to CD4^+^ T cells. Insight into the role of B cell mediated HIV-1 *trans* infection of CD4^+^ T cells is critical to optimizing the effectiveness of HIV-1 therapies in SLO.

**Author summary:** HIV-1 spreads in part by exploiting antigen presenting cells such as B cells, which bind and transfer virus to CD4^+^ T cells through a process called *trans* infection. The molecular events involved in HIV-1 binding to B cells and transfer to CD4^+^ T cells are undefined. B cells frequently interact with CD4^+^ T cells within B cell follicles in secondary lymphoid organs (SLO), which represent a major reservoir of HIV-1. Here, we treated B cells with different SLO-specific activation signals to identify those that facilitate B cell-mediated *trans* infection of CD4^+^ T cells, and the mechanisms involved. We found that B cells stimulated with CD40 ligand (CD40L) and interleukin-4 (IL-4) become highly efficient mediators of *trans* infection due to their increased ability to bind HIV-1. This enhanced binding is driven by upregulation of the C-type lectin receptor CD205, which acts as a critical receptor that facilitates HIV-1 binding and *trans* infection of CD4^+^ T cells. These findings may inform strategies for limiting the viral reservoir in people living with HIV.

## Introduction

While antiretroviral therapy (ART) effectively suppresses HIV-1 replication, it does not eradicate the latent viral reservoir that persists in CD4^+^ T cells [1,2]. Secondary lymphoid organs (SLO), and in particular B cell follicles, represent a major anatomical reservoir of HIV-1 [3–5]. SLO harbor a higher frequency of infected CD4^+^ T cells than peripheral blood due to a combination of factors including virus accumulation on follicular dendritic cells (fDC), enrichment of activated CD4^+^ T cells, chronic inflammation, lack of cytotoxic T-lymphocytes (CTL), and possibly suboptimal drug levels [6–10].

Cell-to-cell transmission of HIV-1, which utilizes the natural interactions among immune cells, is one mechanism of HIV-1 spread [11,12]. Cell-to-cell transmission, or *trans* infection, is 100- to 1000-fold more efficient than direct *cis* infection, likely due to a high multiplicity of infection (MOI) at the site of cell-to-cell contact and is also refractory to inhibition by several classes of ART and neutralizing antibodies [11–14]. *Trans* infection can be mediated by CD4^+^ T cells as well as antigen presenting cells (APC). In the latter, dendritic cells (DC), macrophages, and B cells, bind HIV-1 and subsequently transfer virus to CD4^+^ T cell targets [15,16]. While DC and macrophages can be productively infected via the HIV-1 receptors and coreceptors CD4 and CCR5/CXCR4, respectively, B cells cannot as they do not express CD4 and CCR5 [17].

Various receptors have been implicated in APC-mediated HIV-1 *trans* infection. The C-type lectin receptors DC-specific intercellular adhesion molecule-3 grabbing nonintegrin (DC-SIGN), mannose receptor (MR), and DC-immunoreceptor (DCIR), facilitate HIV-1 *trans* infection by DCs and macrophages [18–21]. Another major receptor, implicated primarily in mature DC-mediated HIV-1 *trans* infection, is the sialic-acid binding immunoglobulin-like lectin 1 (Siglec-1) receptor [22]. In contrast, the receptors involved in B cell-mediated HIV-1 *trans* infection have not been defined.

In SLO, such as lymph nodes, B cells and CD4^+^ T cells constantly interact in and around B cell follicles during the initiation and maintenance of antibody responses with their developments tightly interrelated [23,24]. During these interactions at the border between the B and CD4^+^ T cell areas and in germinal centers (GC), stimulations provided by CD4^+^ T cells can vary depending on the antigen, i.e., interferon-γ (IFN-γ) for a type 1 helper T cell (Th1) response, while other stimulations such as CD40 ligand (CD40L) and interleukin-4 (IL-4), are consistent and are required for B cell survival, proliferation, isotype switching, and affinity maturation [23–25]. Furthermore, B cell survival and maturation in SLO is supported by the B cell activating factor (BAFF), which is secreted by various cells including macrophages, DC, and stromal cells [26]. Due to the frequency of B:T cell interactions and the stimulations provided, B cells in SLO are largely activated, especially during HIV-1 infection [27,28]. The impact of these varying B cell stimulations on HIV-1 binding and *trans* infection remain unknown.

In this study, we sought to determine which B cell stimulations facilitate HIV-1 binding and viral transfer by B cells to CD4^+^ T cells, and to identify the mechanisms involved. We show that, compared to the other stimulations tested, CD40L/IL-4 treated B cells are highly efficient mediators of HIV-1 *trans* infection due to their increased ability to bind virus. Notably, we identified a receptor on B cells, CD205, whose expression is upregulated upon CD40L/IL-4 stimulation, and which facilitates HIV-1 binding, and ultimately *trans* infection of CD4^+^ T cells.

## Results

### CD40L/IL-4 stimulated B cells efficiently *trans* infect CD4^+^ T cells via increased HIV-1 binding

We treated B cells with CD40L/IL-4, CD40L/IFN-γ, BAFF/IL-4, or BAFF/IFN-γ, as described in the Materials and Methods. To assess their capacity to HIV-1 *trans* infect CD4^+^ T cells, we incubated B cells with HIV-1_BaL_ (MOI 10^-3^) at 37°C for 2h, and then co-cultured them with autologous CD4^+^ T cells at a 1:10 ratio of B:CD4^+^ T cell. We found that CD40L/IL-4 stimulated B cells efficiently facilitated HIV-1 *trans* infection of CD4^+^ T cells (Fig 1A). BAFF/IL-4 stimulated B cells also facilitated HIV-1 transmission, but to a significantly lesser extent compared to CD40L/IL-4 (Fig 1A). In contrast, B cells stimulated with IFN-γ (CD40L/IFN-γ or BAFF/IFN-γ) did not support HIV-1 *trans* infection of CD4^+^ T cells (Fig 1A).

**Fig 1.**
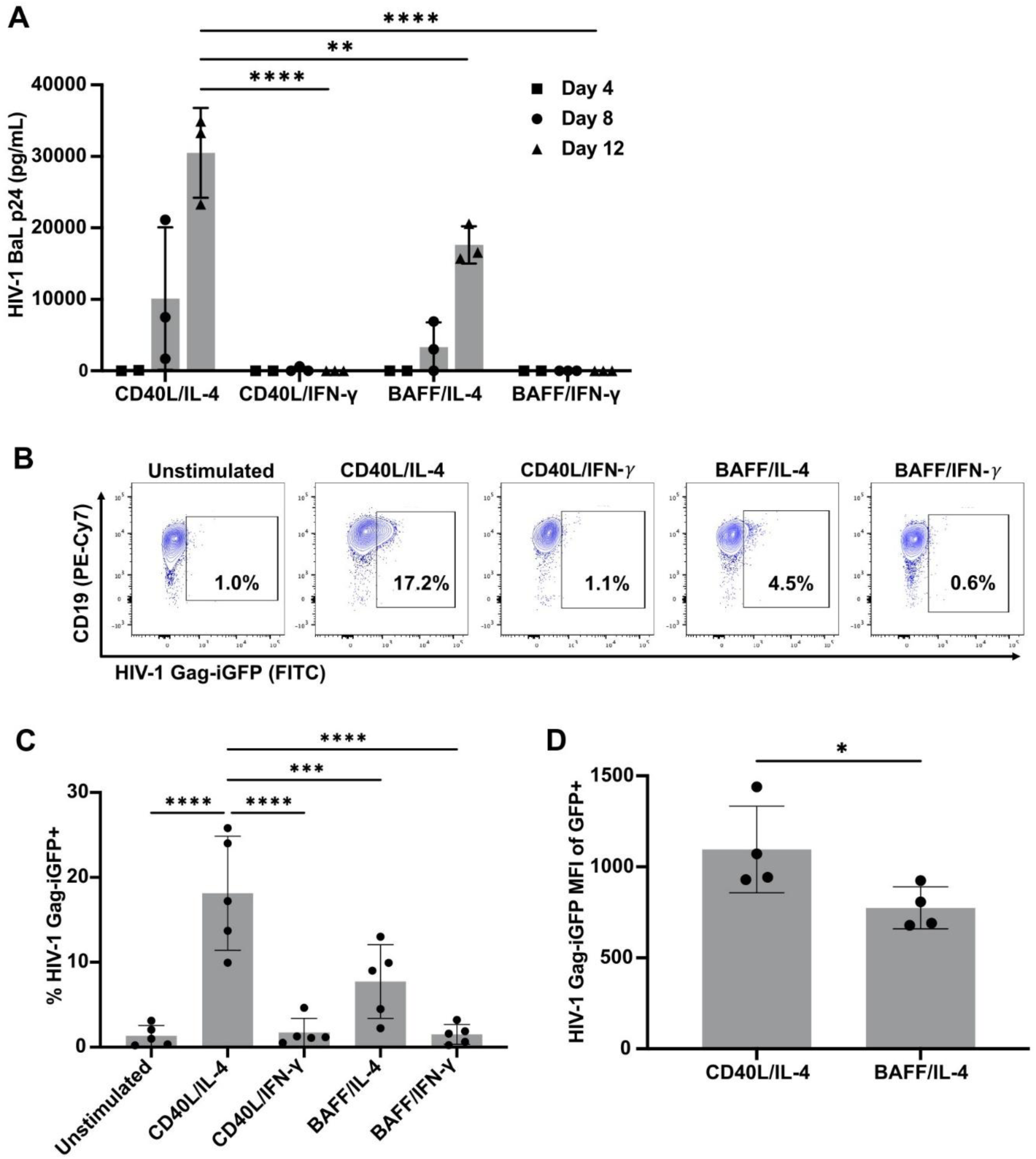
CD40L/IL-4 stimulated B cells enhance HIV-1 binding and *trans* infection of CD4^+^ T cells. **(A)** B cells were stimulated with CD40L/IL-4, CD40L/IFN-*γ*, BAFF/IL-4, or BAFF/IFN-*γ* for 48h. B cells were incubated with HIV-1_BaL_ (MOI 10^-3^) for 2h at 37°C, washed, and co-cultured with CD4^+^ T cells at a 1:10 ratio (B:T cell). Supernatants were collected at days 4, 8, and 12 post co-culture and HIV-1 was quantified by HIV-1 p24 ELISA. Statistical significance was determined by one way ANOVA (p<0.0001) followed by Dunnett’s multiple comparisons for day 12 *trans* infection results; n=3 HIV-1 negative donors; **p<0.005, ****p<0.0001. **(B-D)** B cells were left untreated or stimulated with CD40L/IL-4, CD40L/IFN-*γ*, BAFF/IL-4, or BAFF/IFN-*γ*. B cells were then incubated with HIV-1 Gag-iGFP (MOI 10^-1^) for 2h at 37°C prior to flow cytometry. **(B)** Representative FACS plots. **(C)** Percent of HIV-1 Gag-iGFP positive B cells by stimulation condition. Statistical significance was determined by one way ANOVA (p<0.0001) followed by Dunnett’s multiple comparisons; n=5 HIV-1 negative donors; ***p=0.001, ****p<0.0001. **(D)** HIV-1 Gag-iGFP MFI of the GFP positive population of B cells stimulated with either CD40L/IL-4 or BAFF/IL-4. Statistical significance was determined by the paired t test; n=4 HIV-1 negative donors; *p<0.05.

We next addressed whether this difference in HIV-1 *trans* infection efficiency was due to a difference in the capacity of B cells to bind virus. We evaluated HIV-1 binding by using HIV-1 Gag-iGFP, which has GFP inserted between the matrix (MA) and capsid (CA) Gag domains, making it constitutively fluorescent [29–31]. The extent to which the differentially stimulated B cells bound HIV-1 Gag-iGFP was similar to the *trans* infection data described in Fig 1A. Specifically, CD40L/IL-4 stimulated B cells were the most efficient at binding virus as compared to the other conditions, while BAFF/IL-4 stimulated B cells also bound virus, but to a significantly lesser extent (Fig 1B, C and S1 Fig). In contrast, B cells stimulated with IFN-γ (CD40L/IFN-γ or BAFF/IFN-γ) were largely resistant to HIV-1 Gag-iGFP binding, as were unstimulated B cells (Fig 1B, C). CD40L/IL-4 stimulated B cells also had a significantly higher GFP mean fluorescent intensity (MFI) than BAFF/IL-4 stimulated B cells, suggesting that they also bind more HIV-1 on a per cell basis (Fig 1D). Collectively, these data highlight that the different B cell stimulations impact the ability of B cells to bind HIV-1 and this subsequently results in differences in CD4^+^ T cell *trans* infection efficiency.

### B cell activation alone does not drive HIV-1 binding

We asked whether there was an association between B cell activation and HIV-1 binding. B cell activation was quantified by expression of CD86 and HLA-DR by flow cytometry. Treatment of B cells with CD40L/IL-4, CD40L/IFN-γ, BAFF/IL-4, or BAFF/IFN-γ promoted B cell activation as noted by at least a 2-fold increase in CD86 and HLA-DR expression compared to control, untreated cells (Fig 2A-D). While CD86 expression was significantly upregulated by CD40L/IL-4 stimulation compared to the other conditions, there was not a significant difference in HLA-DR expression between the conditions (Fig 2D). Furthermore, B cells exposed to CD40L/IFN-γ and BAFF/IL-4 displayed similar levels of activation (Fig 2B, D), yet only the BAFF/IL-4 condition supported HIV-1 binding (Fig 1). Collectively, these data suggest that B cell activation alone is not sufficient for HIV-1 binding.

**Fig 2.**
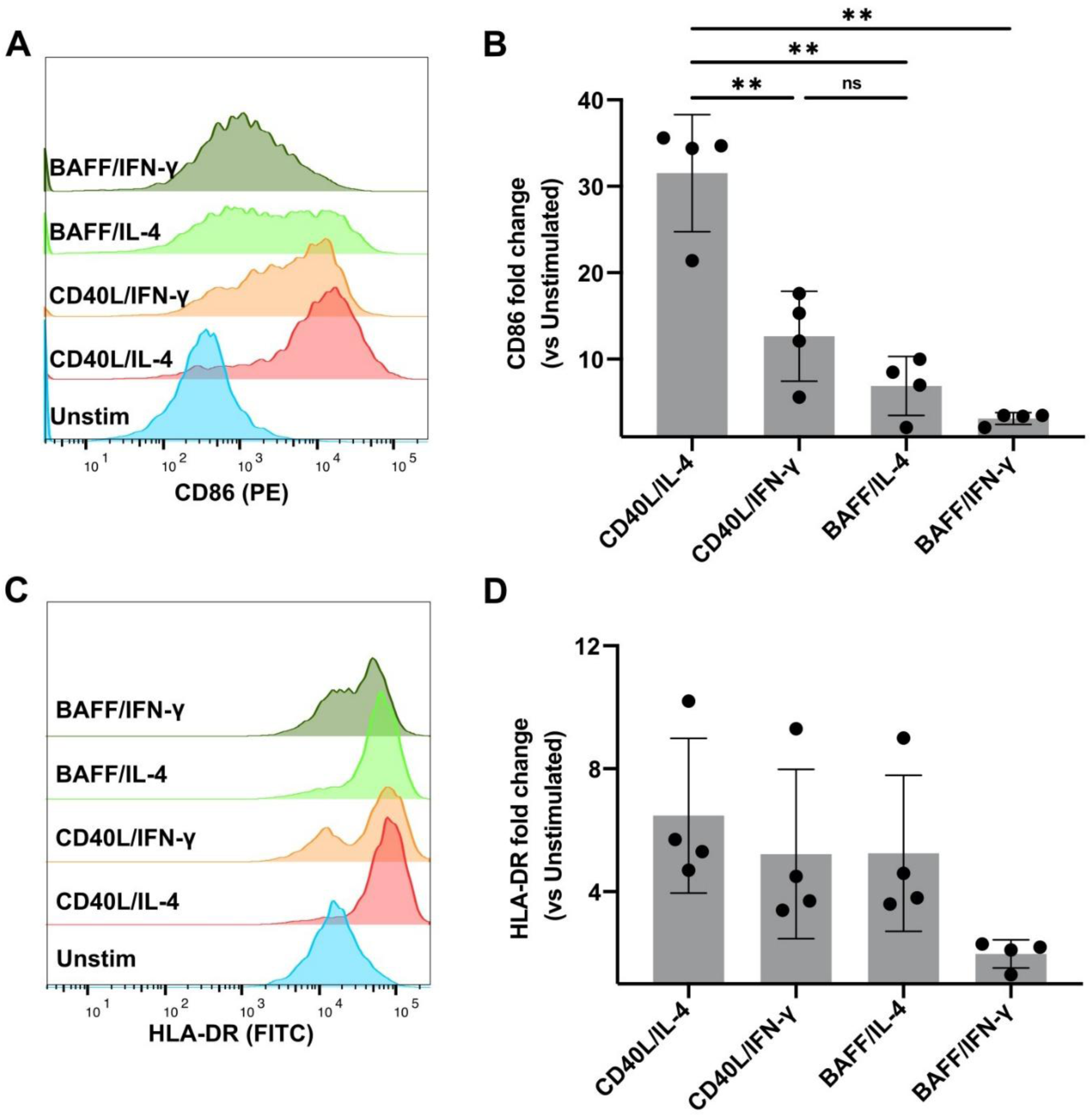
B cell activation alone does not drive HIV-1 binding to B cells. **(A-D)** B cells were left untreated or stimulated with CD40L/IL-4, CD40L/IFN-*γ*, BAFF/IL-4, or BAFF/IFN-*γ* for 48h prior to flow cytometry. Representative histogram of **(A)** CD86 and **(C)** HLA-DR expression. Fold change of **(B)** CD86 and **(D)** HLA-DR MFI expression for each stimulated B cell condition compared to untreated, control B cells. Statistical significance was determined by one way ANOVA (CD86 p<0.0005; HLA-DR p>0.5) followed by Tukey’s multiple comparisons; n=4 HIV-1 negative donors. **p<0.01.

### CD40L and IL-4, but not BAFF, promote HIV-1 binding to B cells

Next, we quantified the ability of B cells exposed to only CD40L, IL-4 or BAFF to bind HIV-1 Gag-iGFP. CD40L treated B cells increased HIV-1 binding in a concentration dependent manner, whereas IL-4 treated B cells increased HIV-1 binding but the effect was not concentration dependent (Fig 3A, B). BAFF treatment alone, did not promote HIV-1 binding to B cells (Fig 3C). These data indicate that both CD40L and IL-4 promote binding of HIV-1 to B cells.

**Fig 3.**
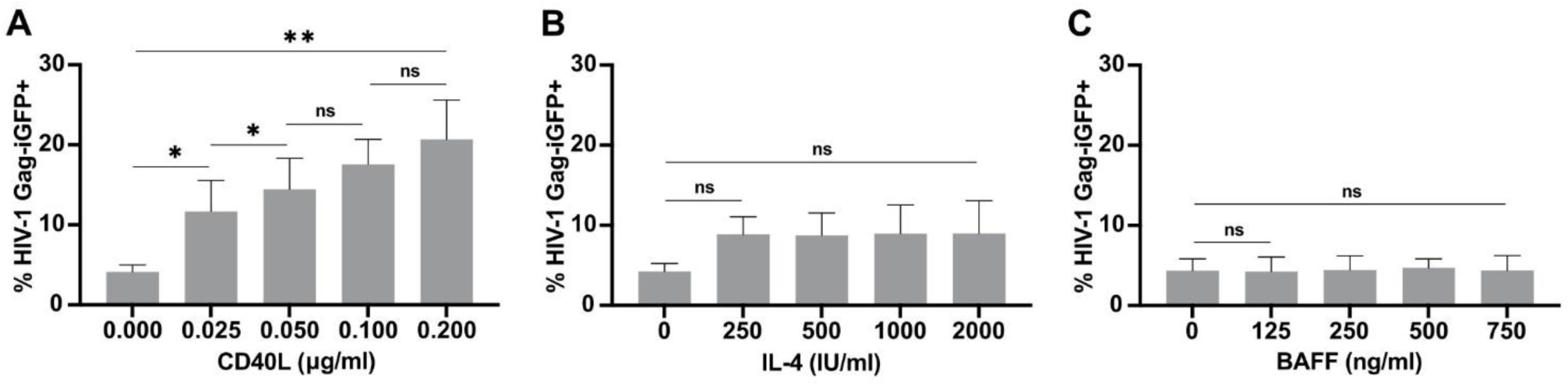
CD40L and IL-4, but not BAFF, promote HIV-1 binding to B cells. **(A-C)** B cells were treated with increasing concentrations of **(A)** CD40L (standard concentration: 0.1 μg/ml), **(B)** IL-4 (standard concentration: 1000 IU/ml), or **(C)** BAFF (standard concentration: 500 ng/ml) for 48h. B cells were then incubated with HIV-1 Gag-iGFP (MOI 10^-1^) for 2h at 37°C prior to flow cytometry. Statistical significance was determined by paired t tests; n=3 HIV-1 negative donors; *p<0.05, **p<0.01.

### CD40L and IL-4 drive expression of CD205 on B cells

To gain insight as to how CD40L/IL-4 stimulation promotes HIV-1 binding to B cells, we performed single cell RNA sequencing (scRNAseq) of B cells from two people without HIV and one person with HIV on ART that were left unstimulated or exposed to CD40L, IL-4, CD40L/IL-4, or CD40L/IFN-γ. Uniform manifold approximation and projection (UMAP) analyses indicated that the B cells exposed to each of these stimulants, or a combination of these stimulants, formed unique clusters based on gene expression (Fig 4A). As noted in the Introduction, the C-type lectin receptors DC-SIGN, MR, and DCIR, and the sialic acid-binding lectin Siglec-1 on DCs and macrophages, are implicated in APC-mediated HIV-1 *trans* infection of CD4^+^ T cells [18–22]. Therefore, we first examined the relative gene expression of each of these receptors on B cells exposed to the different stimulants. We found that there was minimal to no expression of *CD209* (DC-SIGN), *MRC1* (MR), *CLEC4A* (DCIR), or *SIGLEC1* (Siglec-1) on B cells in either the CD40L/IL-4 stimulated condition, or any of the other conditions tested, suggesting that these receptors do not facilitate HIV-1 binding to B cells (Figs 4B and S2).

**Fig 4.**
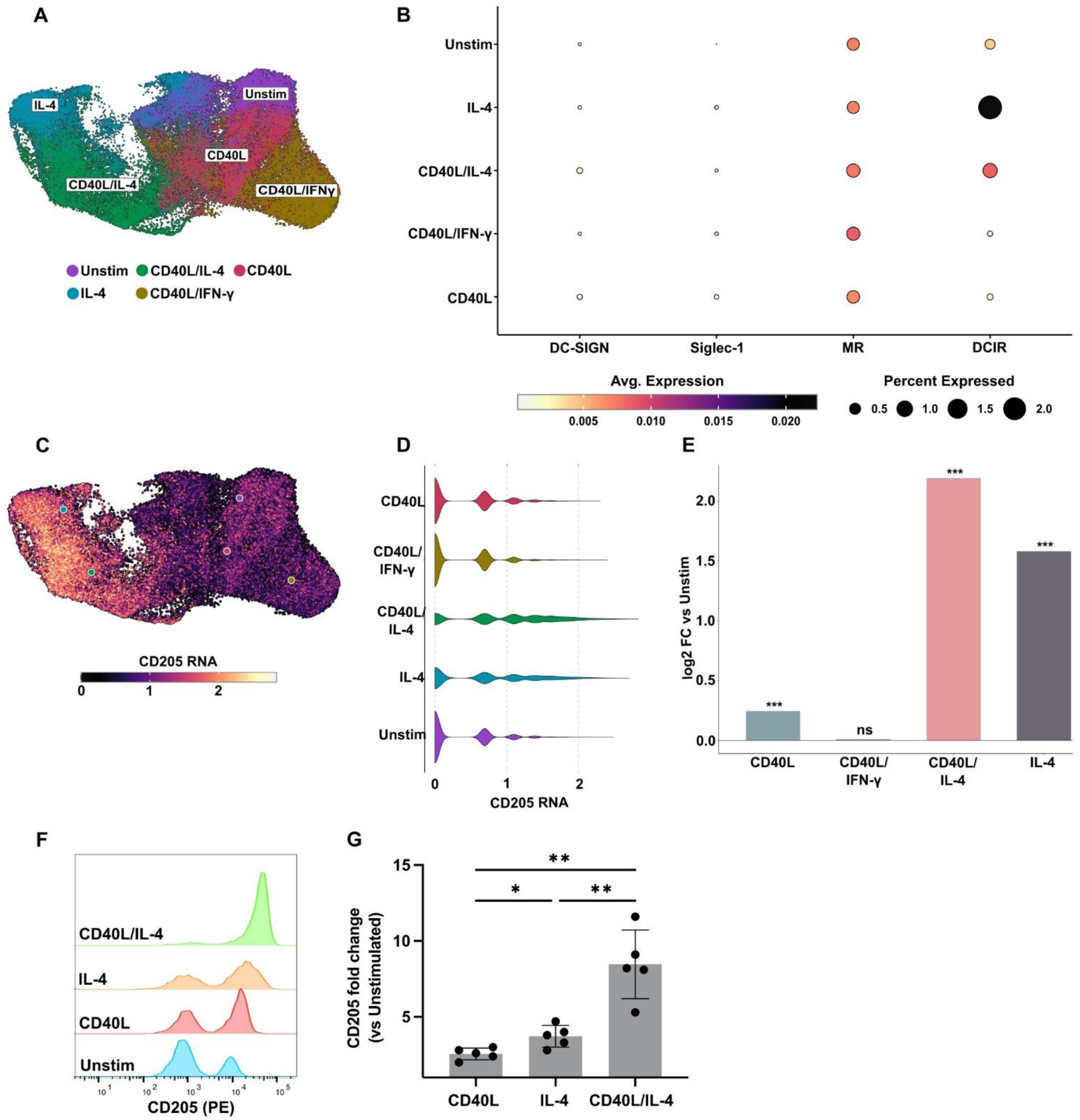
CD40L and IL-4 drive expression of CD205 on B cells. **(A-D)** B cells from 3 donors (2 HIV-1 negative, 1 HIV-1 positive) were left untreated or stimulated with CD40L, IL-4, CD40L/IL-4, or CD40L/IFN-*γ* for 10h prior to scRNAseq. **(A)** UMAP of gene expression of B cells by stimulation condition. **(B)** Dot plot for relative expression of DC-SIGN, Siglec-1, MR, and DCIR represented by average mRNA expression (color) and percent expressed (dot size) for each stimulation condition. **(C, D)** Relative mRNA expression of CD205 by **(C)** UMAP and **(D)** violin plot. **(E)** Bar graph illustrating the fold change in CD205 expression on B cells by condition compared to untreated, control B cells. Statistical significance was determined by MAST; ***p<0.001. **(F, G)** B cells were left untreated or stimulated with CD40L, IL-4, CD40L/IL-4, or CD40L/IFN-*γ* for 48h prior to flow cytometry. **(F)** Representative histogram and **(G)** fold change of CD205 MFI expression for each stimulated B cell condition compared to untreated, control B cells. Statistical significance was determined by one way ANOVA followed by Tukey’s multiple comparisons; n=5 HIV-1 negative donors; *p<0.05, **p<0.01.

Subsequently, we analyzed the mRNA expression levels of all other C-type lectin and Siglec receptors to identify potential HIV-1 binding receptors. Candidates of interest included *LY75*, which encodes the C-type lectin CD205, and FCER2, which encodes the C-type lectin CD23 (Figs S3 and 4C). Further analysis revealed that CD23 did not mediate HIV-1 binding to B cells (data not shown). CD40L/IL-4 stimulation drove the largest increase in CD205 expression on B cells, although both IL-4 and CD40L alone also increased CD205 expression (Figs 4C-4E). When CD205 expression on B cells was examined at the protein level by flow cytometry, we found that both CD40L and IL-4 alone enhanced CD205 expression, but that the combination of the two significantly enhanced its expression (Figs 4F and 4G).

### CD205 mediates HIV-1 binding to CD40L/IL-4 stimulated B cells

We used confocal microscopy to assess whether HIV-1 Gag-iGFP associates with CD205 on B cells exposed to CD40L/IL-4. In addition to CD205, the cells were stained for nuclei (DAPI) and actin (phalloidin) to visualize both the cellular morphology and location of CD205 and HIV-1. We found that HIV-1 Gag-iGFP and CD205 puncta co-localized throughout the B cells (Fig 5A). Z-stack images highlighted similar relative fluorescent intensities of HIV-1 Gag-iGFP and CD205 through the focal depths (Fig 5B). Next, we blocked CD205 on CD40L/IL-4 stimulated B cells using a polyclonal anti-CD205 antibody prior to HIV-1 Gag-iGFP incubation, and then assessed HIV-1 binding as described in Fig. 1B. Antibody-mediated blocking of CD205 significantly decreased HIV-1 binding to B cells (Figs 5C and 5D). This result highlights that CD205 is a critical receptor for B cell-mediated HIV-1 binding.

**Fig 5.**
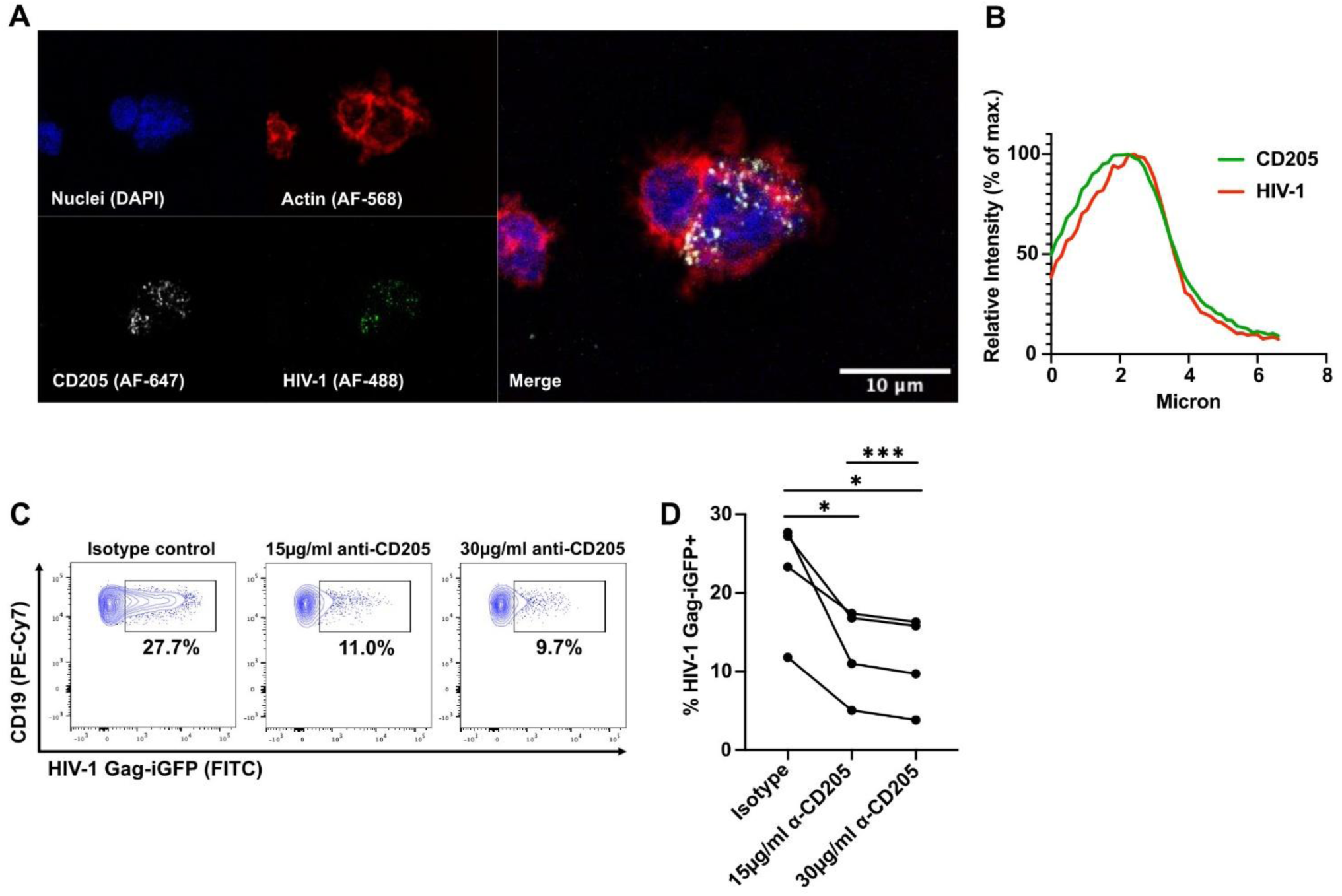
CD40L/IL-4 stimulated B cells bind HIV-1 via CD205. **(A, B)** Confocal microscopy of CD40L/IL-4 stimulated B cells incubated with HIV-1 Gag-iGFP (MOI 10^-1^). Secondary antibody only stained B cells and B – HIV-1 Gag-iGFP samples were used as controls. B cells were stained for nuclei (DAPI), actin (AF-568), and CD205 (AF-647) prior to imaging. **(A)** Confocal images of B cells showing HIV-1 colocalized with CD205. **(B)** Fluorescence intensity plot through the z-axis. Signal intensities were background-subtracted and normalized to the maximum value of each independent channel. **(C, D)** CD40L/IL-4 stimulated B cells were treated with 15 µg/ml isotype control, 15 µg/ml anti-CD205, or 30 µg/ml anti-CD205 polyclonal antibodies for 1h at 37°C prior to incubation with HIV-1 Gag-iGFP (MOI 10^-1^) for 2h at 37° and subsequent flow cytometry. Gates for HIV-1 Gag-iGFP positivity were determined on a B – HIV-1 Gag-iGFP control for each condition. **(C)** Representative FACS plots. **(D)** Percent of HIV-1 Gag-iGFP positive B cells by stimulation condition. Statistical significance was determined by paired t tests; n=4 HIV-1 negative donors; *p<0.05, ***p=0.0004

## Discussion

B cell follicles, in particular germinal centers, serve as a primary anatomical reservoir for HIV-1 infection [3–6]. In untreated people with HIV, the frequency of virus-producing cells in B cell follicles is significantly higher than in extrafollicular areas. In people with HIV on ART, these follicles continue to serve as sanctuary sites where HIV persists. In germinal centers, CD4^+^ T follicular helper and B cells undergo a continuous, bidirectional dialog that drives affinity maturation and directs B cell differentiation into long-lived plasma cells and memory B cells [32–37]. Our group previously demonstrated that, although B cells cannot be productively infected by HIV-1, they can bind HIV-1 and transfer the sequestered virus to uninfected CD4^+^ T cells across a virological synapse, driving highly efficient viral replication [38]. We also reported that B cells, but not other APCs (e.g. DCs), efficiently *trans* infect naive CD4^+^ T cells, thus promoting the establishment and persistence of latent HIV-1 in the resting CD4^+^ T cell compartment [39].

Previous studies reported that DC-SIGN, MR, DCIR, and Siglec-1 on macrophages and DCs support HIV-1 binding and *trans* infection of CD4^+^ T cells [18–22]. However, we found that there was minimal to no expression of these receptors on B cells (Fig 4B and S2). Instead, we identified the C-type lectin CD205 as a B cell receptor that facilitates HIV-1 binding. Prior research also reported that CD205 on human kidney tubular (HK2) cells acts as an HIV-1 receptor [40,41]. Similar to B cells, renal tubular cells do not express any of the known HIV-1 receptors (CD4, CCR5, CXCR4, DC-SIGN or mannose receptors), and interaction of HIV-1 with CD205 on the HK2 cells resulted in internalization of the virus and establishment of a non-productive infection. However, HIV-1 could be transmitted from HK2 cells in *trans* to CD4^+^ T cells.

CD205 is expressed on several cell types including conventional DCs [42]. CD205 is also expressed at moderate levels on B lymphocytes, with CD40L and IL4 exposure further increasing its expression, as reported in this study. Of note, GC B cells also express CD205 [43]. On B cells, CD205 serves as a critical surface receptor that regulates B cell activation, antigen capture, and downstream immune communication. Typically, once a ligand binds to the external C-type lectin domains of CD205, the receptor relies on a specific coated-pit motif in its cytoplasmic tail to trigger rapid clathrin-mediated endocytosis. The internalized antigens are targeted into late endosomes or lysosomes that are rich in MHC II products, where antigen is processed for presentation in the context of MHC II to T cells [44]. How HIV-1 persists in B cells allowing it to be transferred to CD4^+^ T cells is unknown. Either the virus may escape degradation or persist in endosomal compartments. Of note, DC-SIGN mediated internalization of HIV-1 has been reported to stabilize the virus and preserve its infectivity [45].

In conclusion, this study identifies CD205 as a receptor on B cells that facilitates HIV-1 binding and transfer of virus to CD4^+^ T cells. We speculate that in B cell follicles, HIV-1 hijacks the natural interactions between B and CD4^+^ T cells, allowing for highly efficient *trans* infection of CD4^+^ T cells and thus promoting persistent infection and reservoir maintenance. This work has implications for potential cure strategies as it highlights the importance of therapies to target the HIV-1 reservoir not just in the peripheral blood, but also in tissue, and specifically, the B cell follicles.

## Materials and Methods

### B and CD4^+^ T cell isolation and culture

Peripheral blood mononuclear cells (PBMC) were isolated from buffy coat blood from Vitalant (Pittsburgh Blood Bank) or from whole blood from the University of Pittsburgh Clinical Research Site of the MACS/WIHS Combined Cohort Study (MWCCS) through standard density gradient separation using lymphocyte separation medium (Corning). All participants signed consent forms allowing the blood products to be used for research and IRB approval (CR19110166-007) was attained for working with human blood samples. B and CD4^+^ T cells were purified from PBMCs using the following separation kits: human CD4 MicroBead positive selection (Miltenyi Biotec) and EasySep human B cell negative selection (STEMCELL). All kits were used per the manufacturer’s instructions. CD4^+^ T cells were cultured for two days in 6-well plates at a concentration of 2×10^6^/ml with CCL19 (100 nM; R&D Systems) [46]. B cells were cultured for two days in 24 well plates at a concentration of 1×10^6^/ml with one or a combination of the following stimulations: CD40L (MEGACD40L Protein, recombinant human; 0.1 μg/ml; Enzo Life Sciences), IL-4 (recombinant human; 1,000 U/ml; R&D Systems), IFN-γ (recombinant human; 1,000 U/ml; R&D Systems), and BAFF (recombinant human; 500 ng/ml; Peprotech). CD4^+^ T and B cells were cultured in complete media of Iscove’s Modified Dulbecco’s Medium (IMDM) (Gibco) containing 10% human serum AB (Gemini Bio), gentamicin (0.5%; Gibco), and GlutaMAX (1%; Gibco).

#### *Trans* infection assay

CCR5-tropic HIV-1_BaL_ was purified from PM1 cells [47]. B cells were incubated with HIV-1_BaL_ (MOI 10^-3^) at 37°C for 2h, washed 3 times with media, and subsequently co-cultured with autologous CD4^+^ T cells at a 1:10 ratio (B: CD4^+^ T cell) in complete media in 96-well round bottom plates. As controls, *cis* infections for each donor were also included in which CD4^+^ T cells were incubated with HIV-1_BaL_ at MOI 10^-3^ (*cis* low) and MOI 10^-1^ (*cis* high) at 37°C for 2h, washed 3 times with media, and cultured alone. Supernatants were collected at days 4, 8, and 12 post co-culture. Infection was measured by HIV-1 p24 antigen capture via enzyme-linked immunosorbent assay (ELISA) (Leidos Biomedical Research, Frederick National Laboratory for Cancer Research) per the manufacturer’s instructions.

### Flow cytometry

The LIVE/DEAD fixable aqua or violet dead cell stains (Life Technologies) were used for viability exclusion and the following antibodies were used for immunostaining: CD19-PE-Cy7 (clone HIB19, BioLegend), CD20-PerCP-Cy5.5 (clone 2H7, Invitrogen), CD205-PE (clone MG38, BD Pharmingen), CD80-FITC (clone MAB104, Beckman Coulter), CD86-PE (clone HA5.2B7, Beckman Coulter), and HLA-DR-FITC (clone TU36, BD Pharmingen). Staining was performed in FACS buffer composed of 1x PBS (Cytiva), bovine serum albumin (BSA) (0.5%; Sigma Aldrich), HEPES (10 mM; Gibco), and EDTA (5 mM; Life Technologies Invitrogen). Samples were fixed using 2% PFA and subsequently stored in FACS buffer. Samples were analyzed on a BD LSRFortessa flow cytometer. Gates and relative expression were based on fluorescence minus one controls, unstained controls, or unstimulated samples. Data were analyzed using FlowJo version 10.10.0 (BD Biosciences).

### HIV-1 binding assay

The plasmid for HIV-1 Gag-iGFP_JRFL was obtained from BEI Resources donated by Dr. Benjamin Chen (Cat# 12456). This plasmid was used to generate fluorescently labeled CCR5-tropic HIV-1 (referred to as HIV-1 Gag-iGFP). Isolated plasmid DNA was transfected into HEK293T cells cultured in DMEM (Corning) with 10% fetal bovine serum (FBS) (Optima) using opti-MEM (Gibco) and lipofectamine 2000 (Invitrogen). Two days post transfection, supernatants were collected and virus was concentrated using Amicon ultra centrifugal filters (100kDa; Millipore). Virus infectivity and infectious units/ml were determined by infecting TZM-bl cells with dilutions of HIV-1 Gag-iGFP and HIV-1_BaL_ (known titer) using the britelite plus (Revvity) luciferase reporter gene assay system. Luminescence was measured by the Vario Skan Lux plate reader (Thermo Scientific). For HIV-1 binding flow cytometry assays, B cells were incubated with HIV-1 Gag-iGFP (MOI 10^-1^) at 37°C for 2 hours in 96-well V-bottom plates and then washed to remove excess virus. The cells were stained, fixed, and analyzed on an LSR Fortessa flow cytometer. To measure HIV-1 binding, expression in the FITC channel was used due to its similar emission spectrum to GFP. HIV-1 Gag-iGFP positivity on samples of B cells + HIV-1 Gag-iGFP were determined using gates set on controls of B cells – HIV-1 Gag-iGFP. For CD205 blocking assays, B cells were incubated with human BD FC block (2.5 μg; BD) at room temperature for 10 minutes. The cells were then incubated with 15 μg/ml or 30 μg/ml of human CD205 antibody (polyclonal goat IgG; R&D Systems) or normal goat IgG control (polyclonal goat IgG; R&D Systems) at 37°C for 1 hour prior to HIV-1 Gag-iGFP incubation.

### Single cell RNA sequencing (scRNAseq)

B cells from 2 MWCCS participants (1 HIV-1 positive, 1 HIV-1 negative) and 1 blood bank donor (HIV-1 negative) were left unstimulated or stimulated with CD40L, IL-4, CD40L/IL-4 or CD40L/IFN-γ. Stimulation conditions were multiplexed using BD Rhapsody™ Single Cell Multiplexing Kit. Samples were then mixed in equal parts before loading onto a BD Rhapsody™ 8-Lane Cartridge per the manufacturer’s instructions. Single cell capture was performed using the BD Rhapsody™ HT Xpress Single-Cell Analysis System, and quality control metrics were verified using the BD Rhapsody™ Scanner. Whole transcriptome cDNA libraries were generated according to the manufacturer’s instructions and submitted to the UCLA Technology Center for Genomics and Bioinformatics, and sequencing data was acquired on an Illumina NovaSeq X. Initial cell calling, quality filtering, alignment, and annotation were performed using the BD Rhapsody™ WTA Analysis Pipeline on the Seven Bridges Genomics Platform. The data were then imported into R Studio version 4.5.2, and the Seurat package version 4.3.0 was used to remove multiplets and cells with mitochondrial reads > 25%. The Seurat package was also used to integrate data from the 3 participants for combined analysis. Single, live cells (n=67,542) were analyzed and assessed for differential gene expression. The SCpubr packages were used to generate uniform manifold approximation projections (UMAP), violin plots, bar graphs, and heat maps.

### Confocal microscopy

B cells were adhered to slides using Cell-Tak cell tissue adhesive (Corning) before being fixed in 4% PFA (Thermo Scientific) in a sucrose medium (30mM sucrose (Sigma) + 1x PBS (Cytiva)) at 37°C for 15 minutes. Slides were washed with PBS in Coplin jars and stored in sucrose media at 4°C overnight. Slides were washed in PBS and permeabilized using 0.1% triton (Sigma Aldrich) for 15 minutes at room temperature.

They were washed 3 times with PBS followed by 3 times with PBB (1x PBS + 0.5% BSA). The slides were blocked with 10% normal goat serum (Invitrogen) in PBS at room temperature for 30 minutes followed by 2 washes with PBB. The primary CD205 polyclonal antibody (rabbit anti-human; Invitrogen) was added at a concentration of 1:100 in PBB to all slides except for the secondary only control for 1 hour at room temperature followed by 5 washes with PBB. The secondary CD205 antibody (goat anti-rabbit IgG-AF-647; Invitrogen) was added at a concentration of 1:500 in PBB and the secondary GFP polyclonal antibody (rabbit IgG anti-GFP-AF-488; Invitrogen) was added at a concentration of 1:200 in PBB for 1 hour at room temperature followed by 5 washes with PBB. To stain actin, phalloidin-AF568 (Invitrogen) was added at a concentration of 1:200 in PBB. At the same time, 4’,6-diamidino-2-phenylindole (DAPI) (Thermo Scientific) was added at a concentration of 1:500 in PBB. Phalloidin and DAPI were incubated on the slides for 30 minutes at room temperature followed by 2 washes with PBB, 2 washes with PBS, and 1 wash with distilled water. Coverslips were mounted on slides with ProLong Glass Antifade Mountant (Invitrogen) and left to dry at room temperature overnight. Slides were imaged at the University of Pittsburgh Center for Biologic Imaging (CBI) using Nikon A1 microscopes. Analysis of images used Fiji Image J (Version 2.16.0) and NIS Elements Viewer using maximal intensity projections (MIP) of Z-stacks.

### Statistics

HIV-1 binding and *trans* infection data were analyzed using GraphPad Prism version 10.6.1. Normality was determined by the Shapiro-Wilk test. The one-way ANOVA was used to measure the significance of differences between the means of three or more groups followed by Dunnett’s or Tukey’s multiple comparisons tests. The paired t test was used to determine statistical significance between two related groups.

## Supplementary Figure Legends

**S1 Fig.**
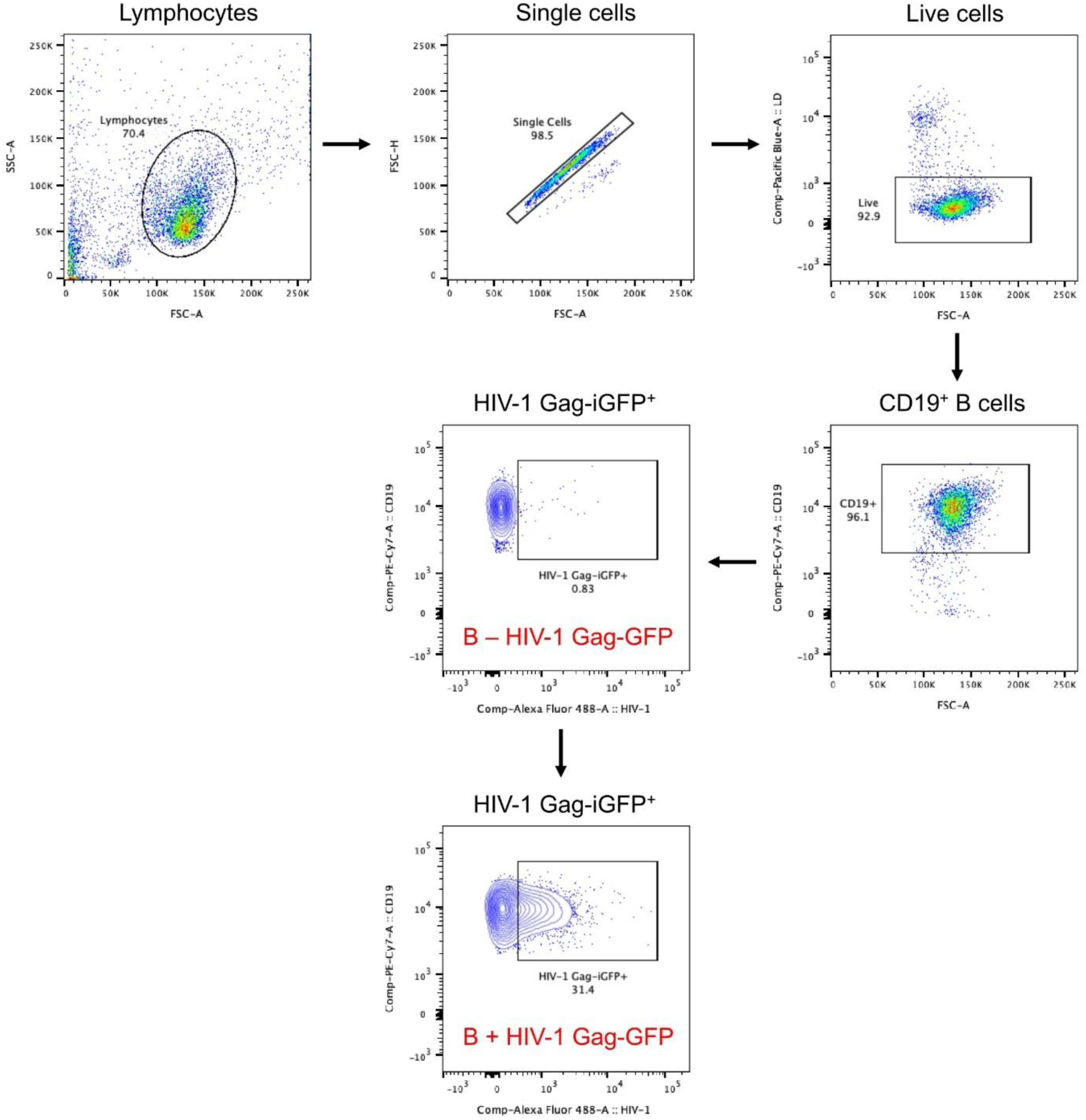
Gating strategy for determining the percentage of HIV-1 Gag-iGFP positive B cells. Gating was performed by forward and side scatter and on single and live cells. CD19 expression was used as a B cell marker. Gates for HIV-1 Gag-iGFP positivity were determined on a B – HIV-1 Gag-iGFP control for each stimulation or experimental condition. Representative example of a B – HIV-1 Gag-iGFP control used for determining HIV-1 Gag-iGFP positivity in a B + HIV-1 Gag-iGFP condition.

**S2 Fig.**
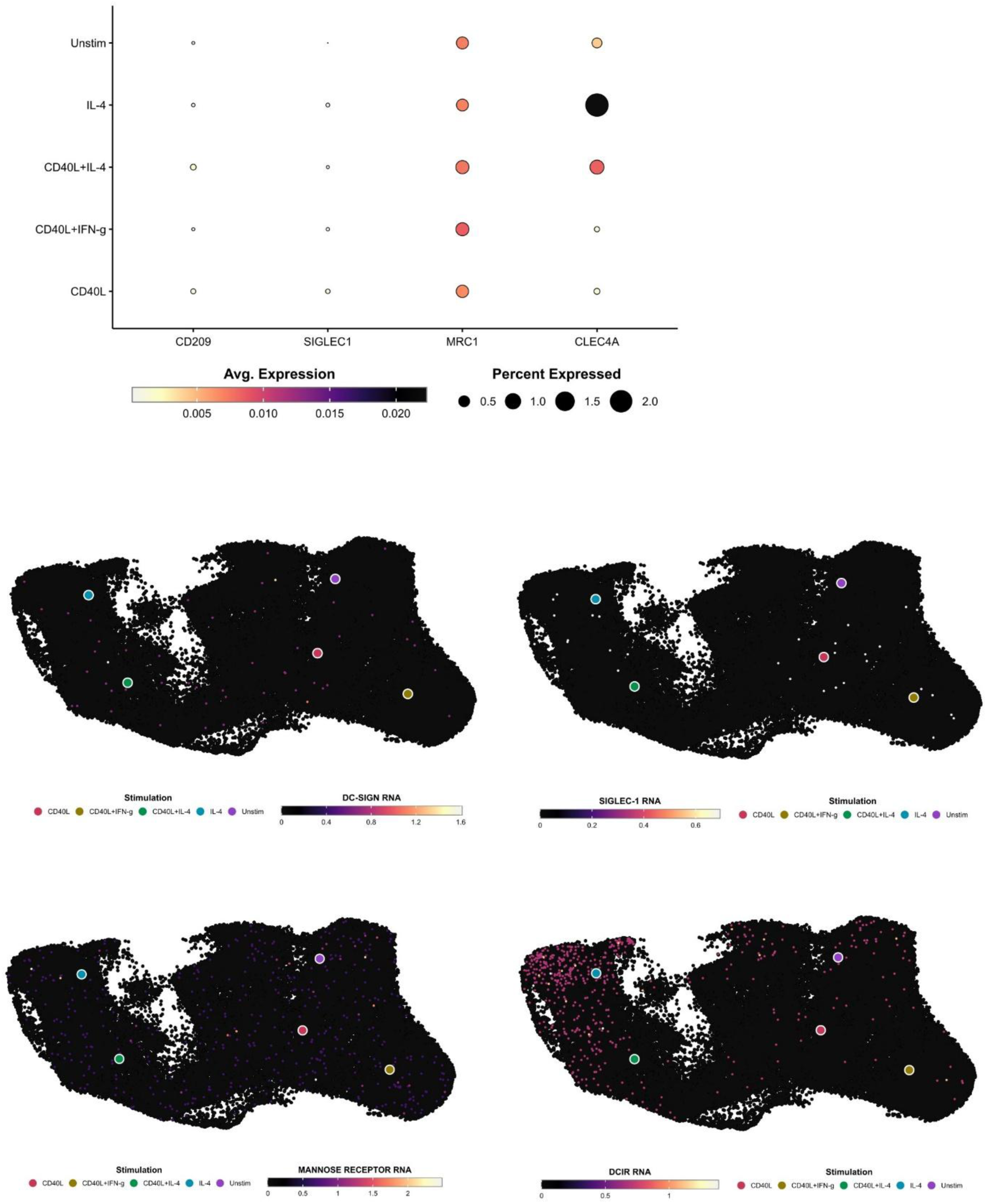
Expression of HIV-1 *trans* infection receptors by B cells. UMAPs of relative mRNA expression of *CD209* (DC-SIGN), *SIGLEC1* (Siglec-1), *MRC1* (MR), and *CLEC4A* (DCIR) for each stimulation condition.

**S3 Fig.**
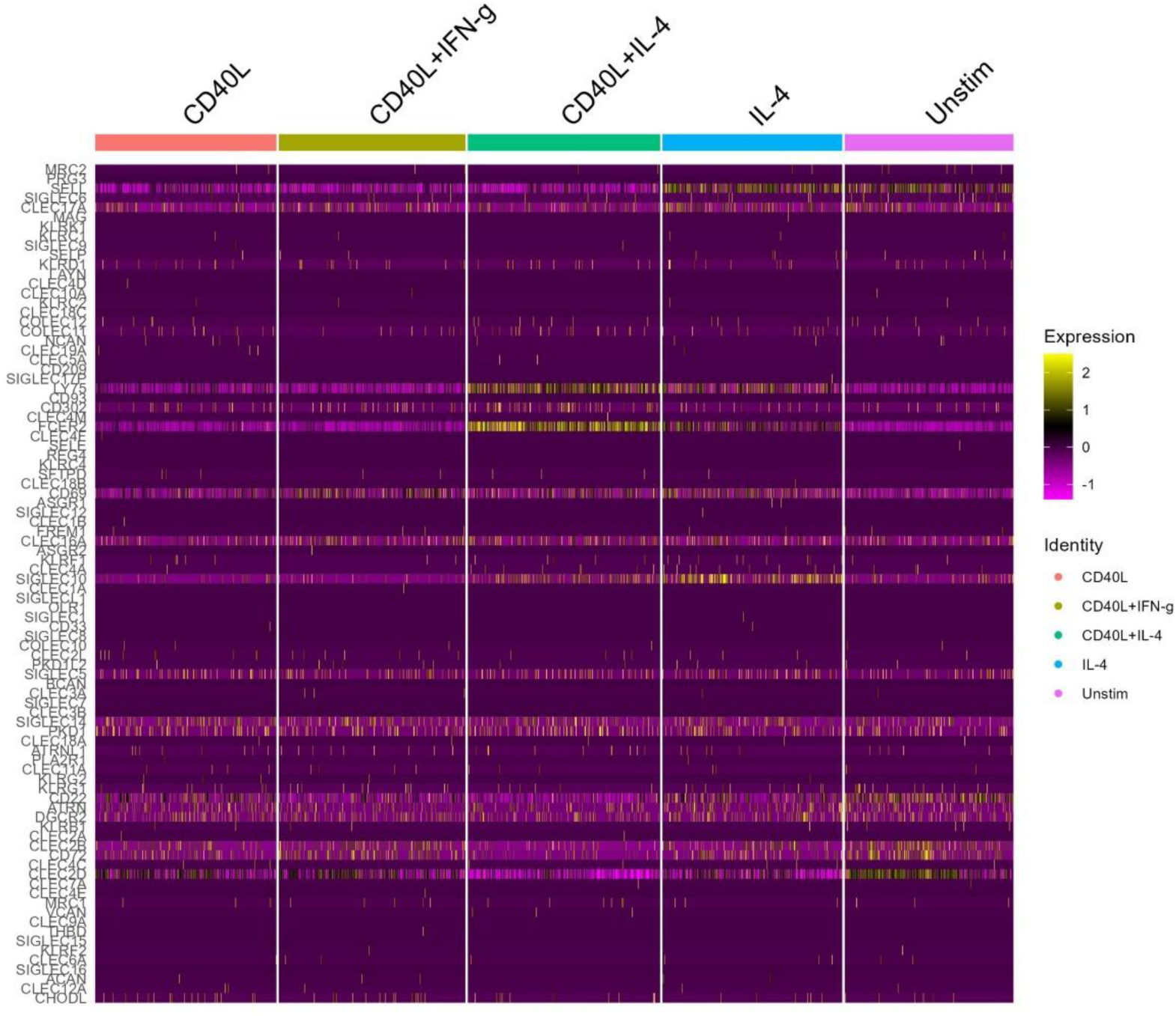
Expression of C-type lectin and Siglec receptors by B cells. Heat map of relative mRNA expression by each cell of all C-type lectin and Siglec receptors separated by stimulation condition.

## Acknowledgments

The research was supported by the following funding from the National Institutes of Health: R01 AI162615, U01 HL14620801, 1S10OD040139-01 and the Rustbelt CFAR (2P30 AI036219). The contents of this publication are solely the responsibility of the authors and do not represent the official views of the National Institutes of Health (NIH). The authors gratefully acknowledge the contributions of the study participants and dedication of the staff at the Pittsburgh MWCCS CRS site. The authors thank Holly Bilben, Lori Caruso, Kathy Hartle, Kathy Kulka, Patrick Mehta, Giovanna Rappocciolo, and the University of Pittsburgh Center for Biologic Imaging for technical assistance.

